# DeepPaint: A deep-learning package for Cell Painting Image Classification

**DOI:** 10.1101/2024.10.08.617198

**Authors:** Diego Luna, Erik C. Johnson, Laura J. Dunphy, Ryan McQuillen

**Author notes:** Funding Statement: This research was supported by internal funding from the Johns Hopkins University Applied Physics Laboratory.

## Abstract

Recent developments in the high-content imaging (HCI) space have allowed for the production of large Cell Painting datasets. These datasets are typically derived from cells exposed to a set of biological perturbants including proteins, small molecules, or even pathogens. While the method of Cell Painting has shown utility for drug discovery and hazard evaluation purposes, traditional analyses pipelines applied Cell Painting datasets typically require the segmentation of single cells from thousands to millions of images, a process that is time-consuming and subject to noise and experimental variability. Here we present DeepPaint, a Python package that uses a deep learning framework to perform image analysis of cell painting images including treatment classification and latent space analysis, circumventing the need for image segmentation. DeepPaint is easily tunable to different HCI setups and datasets and can be applied to classify broad types of biological perturbations. Here we demonstrate that DeepPaint can generate highly accurate neural networks for binary and multiclass classification of cell painting images. The DeepPaint package and example notebooks are freely available at https://github.com/jhuapl-bio/DeepPaint.

## INTRODUCTION

Image-based morphological profiling (IBMP) assays that employ the method of Cell Painting have emerged as promising high-throughput screening methods for drug discovery and chemical characterization^1–5^. In a typical Cell Painting-based profiling assay^6,7^, cells are subjected to a large library of biological perturbants including small molecules or protein factors^1–4,8,9^. Following exposure, the cells are fixed, stained, and imaged in a high-throughput manner^6,7^. Traditionally, the multi-channel cellular images are segmented down to the single-cell level, from which hundreds to thousands of measurements per cell are derived and used to construct a cellular “phenoprint” using software such as the CellProfiler package^10–12^. Phenoprints are then typically aggregated across wells or conditions and compared across perturbants.

While this method has proven generally successful for relating bioactive compounds by mechanism^1,8,9,13^ and for tying phenotypic data with orthogonal genetic and proteomic assays^5,14–16^, cell segmentation-based methods have several downfalls including high computational power requirements, slow segmentation speeds, and inaccuracies in defining cell boundaries^17^. Additionally, the *a priori* definition of image-based measurements can, in some cases, limit the statistical power of downstream analysis by capturing extraneous noisy measurements that do not add useable information^5,9,17^. To add to these difficulties, experimental variation in individual profiling datasets can impede the ability to compare multiple experiments from different experimental sources^18^.

To circumvent these issues, novel deep-learning-based methods that interpret and classify whole cell images have begun to be developed that exclude the need for *a priori* measurement definition that can impede analytical performance^1,17,19^. Here we present a new Python package, DeepPaint, a DenseNet-based IBMP image classification pipeline designed for use with Cell Painting datasets (**Figure *1***). DeepPaint is a user-friendly package that requires only a directory of images, metafile of image labels, and a user-updated YAML file as inputs and can be used to easily generate neural network-based binary and multiclass classifiers from Cell Painting data. Notably, DeepPaint has been designed specifically to handle the nuances of Cell Painting datasets, including those of varying image sizes and channel numbers. Additionally, DeepPaint has built-in data normalization and data augmentations methods that can be applied during model training to help improve model performance and reduce the impact of experiment-to-experiment variability. Lastly, during inference, DeepPaint outputs quantitative performance metrics as well as data visualizations of the embedding latent space to enable rapid comparison across experimental conditions. Altogether, this makes DeepPaint a user-friendly, publicly available tool that enables discovery and classification of large datasets of Cell Painting images for a broad range of users.

**Figure 1.**
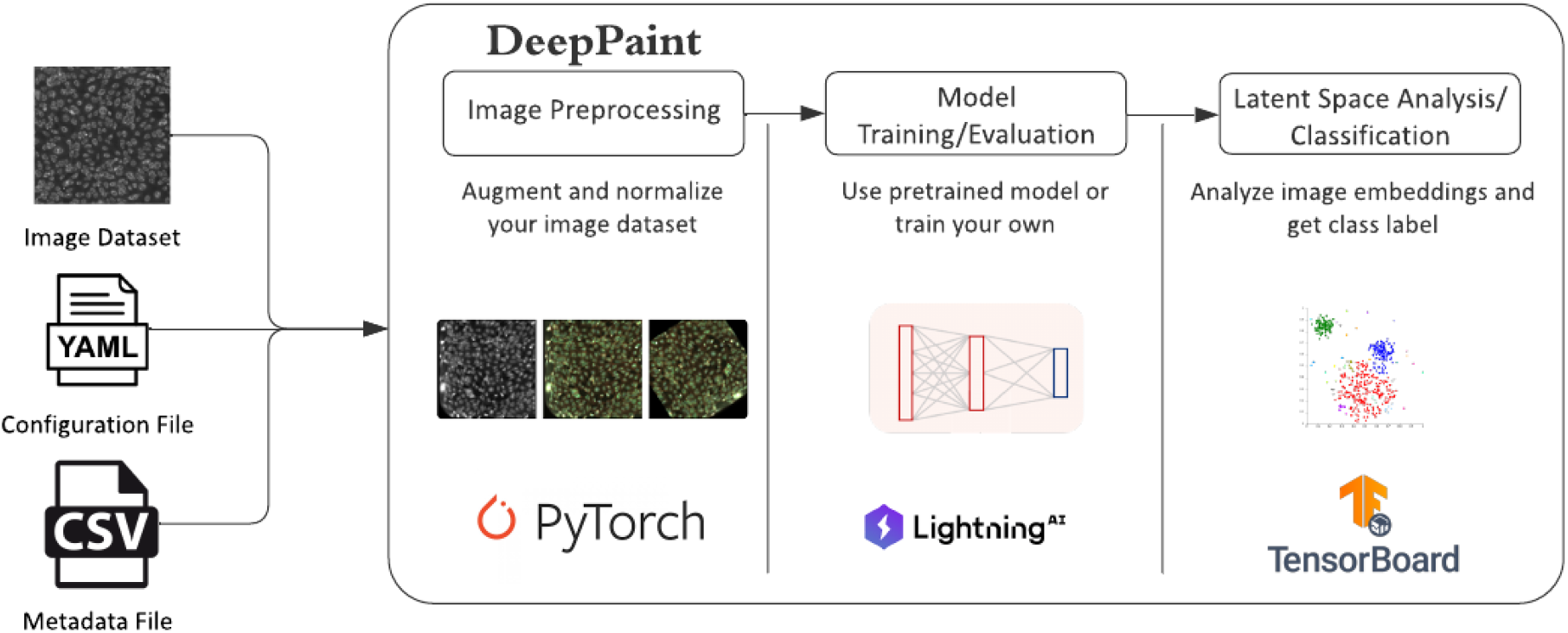
Overview of the DeepPaint package. This package uses a DenseNet model to classify Cell Painting images and compare image embeddings.

## Methods

### DeepPaint framework

DeepPaint is a Python-based^20^ package for the classification of multi-channel cellular images designed to work with Cell Painting^6,7^ datasets. This package relies on PyTorch^21^ Lightning^22^, an open-source deep-learning framework that we used to implement a DenseNet-161 densely connected convolutional neural network architecture^23^ for image classification. The utility of the DenseNet-161 architecture to interrogate Cell Painting data has been demonstrated by the private sector^1^; however, our network is distinct from previous efforts^24^ in that we modified the initial layer of the standard DenseNet-161 to work directly with multi-channel Cell Painting images rather than reducing the multi-channel images down to 3-channel RGB images prior to network training. Additionally, DeepPaint, provides 1) a number of preprocessing features to account for image and experimental variability, 2) a package for latent space analysis of features from the DenseNet and 3) network weights for existing datasets. For this work, experimental hyperparameters are described in the configuration files of the DeepPaint repository to aid in reproducibility.

### DeepPaint implementation

DeepPaint provides a user-friendly framework with a single driver script as the entry point for training and inference, and a complementary script for downstream generation of image embeddings. Both scripts are used in conjunction with a YAML file containing all the necessary user-defined parameters for training and inference. The DeepPaint configuration allows for specification of standard machine learning parameters, such as gradient estimation methods, learning rates, and learning rate schedulers. It enables data selection, finetuning and usage of pretrained network weights. Additionally, the scripts provide specific cell painting functionality:

- Preprocessing of multi-channel cell painting images
- Generation of embeddings of cell painting objects
- Generation of key plots for analysis

The driver script provided can fulfill different use cases. For instance, inference with pre-trained networks can be used to analyze unknown samples in the cell painting latent space or perform high-throughput screening. The driver script supports training given new data, but also fine tuning from existing checkpoints, allowing for different configurations of frozen layers.

### Training and inference on a public cell painting dataset

The public dataset, RxRx2, generated by the company Recursion^25^ was used to develop and evaluate our DeepPaint algorithm. This dataset consists of 131,953 images of HUVEC cells treated with over 400 unique compounds at a 6-point dose range. Cytokines, growth factors (GFs), and toxins shown by Cuccarese, *et. al*., to have elicited a response in HUVEC cells along with untreated controls were selected from RxRx2 training with DeepPaint. This resulted in a final set of 37 compounds and untreated controls.

For binary classification, images were labeled as either “low dose” (*e*.*g*., untreated and lowest three tested concentrations of each compound) or “high dose” (*e*.*g*., highest three tested concentrations of each compound). For the multi-class problem, the lowest three tested concentrations were excluded from the training and validation data and images were labeled as untreated, cytokine/GF, or toxin, where cytokine/GF and toxin contained images of cells exposed the highest three tested concentrations of each compound. The size of image datasets used for training and inference of each classification problem are summarized in **Table 1**. Image embeddings, true and predicted class membership, and associated metadata are provided in the supplement for the reported binary (**Data S1-S3**) and multiclass classifiers (**Data S4-S6**).

**Table 1.** Summary of training and inference datasets.

| Label | Class | # Training Images | # Validation Images | # Test Images |
| --- | --- | --- | --- | --- |
| *Binary* |  |  |  |  |
| 0 | Low Dose | 9028 | 1935 | 1935 |
| 1 | High Dose | 3716 | 796 | 797 |
| *Multiclass* |  |  |  |  |
| 0 | Cytokine-GF | 2617 | 561 | 561 |
| 1 | Toxin | 1099 | 235 | 236 |
| 2 | Untreated | 5305 | 1137 | 1137 |

### Data and Software availability

The DeepPaint Python package is publicly available for download. Installation and usage instructions as well as all metadata used in this analysis can be found at https://github.com/jhuapl-bio/DeepPaint. Public images used in this study can be downloaded directly from: https://www.rxrx.ai/rxrx2.

## Results

### Image Normalization and Binary Classification Performance

To evaluate the performance of DeepPaint, we used a subset of a publicly available dataset (RxRx2) containing two replicate experiments (HUVEC-1 & HUVEC-2) of HUVEC cells exposed to a variety of protein and small molecule perturbants over a log dose range spanning 6 orders of magnitude in units of ng/mL^26^. For our first experiment, we used this data to create a binary classification problem where images corresponding to rank-order does between 0 and 3 were labeled as “low dose” or and those from rank order 3 to 6 were labeled as “high dose”. Notably, our labeling scheme resulted in an imbalanced dataset (**Table *1***) as untreated control cells were contained within the “low dose” population.

Prior to training we implemented an image normalization strategy to minimize the variability typically seen in Cell Painting datasets^18,24^. For training we used 70% of our labeled images to train our DenseNet-161 model for 100 epochs starting with random weights (**Error! Reference source not found**.). Visualization of our model output using principal component analysis (PCA) colored by experiment showed a large degree of overlap between the data each replicate experiment (**Figure 2A**) suggesting that our upstream image normalization pipeline was able to effectively minimize variability in the input experimental data. Importantly, when we colored our model output by treatment class (**Figure 2B**) we saw a high degree of separation accompanied by excellent model statics (**Figure 2C-D, F1 score: 0.85**, **precision: 0.82, recall: 0.89**) suggesting our model was viable for more complex image classification tasks.

**Figure 2.**
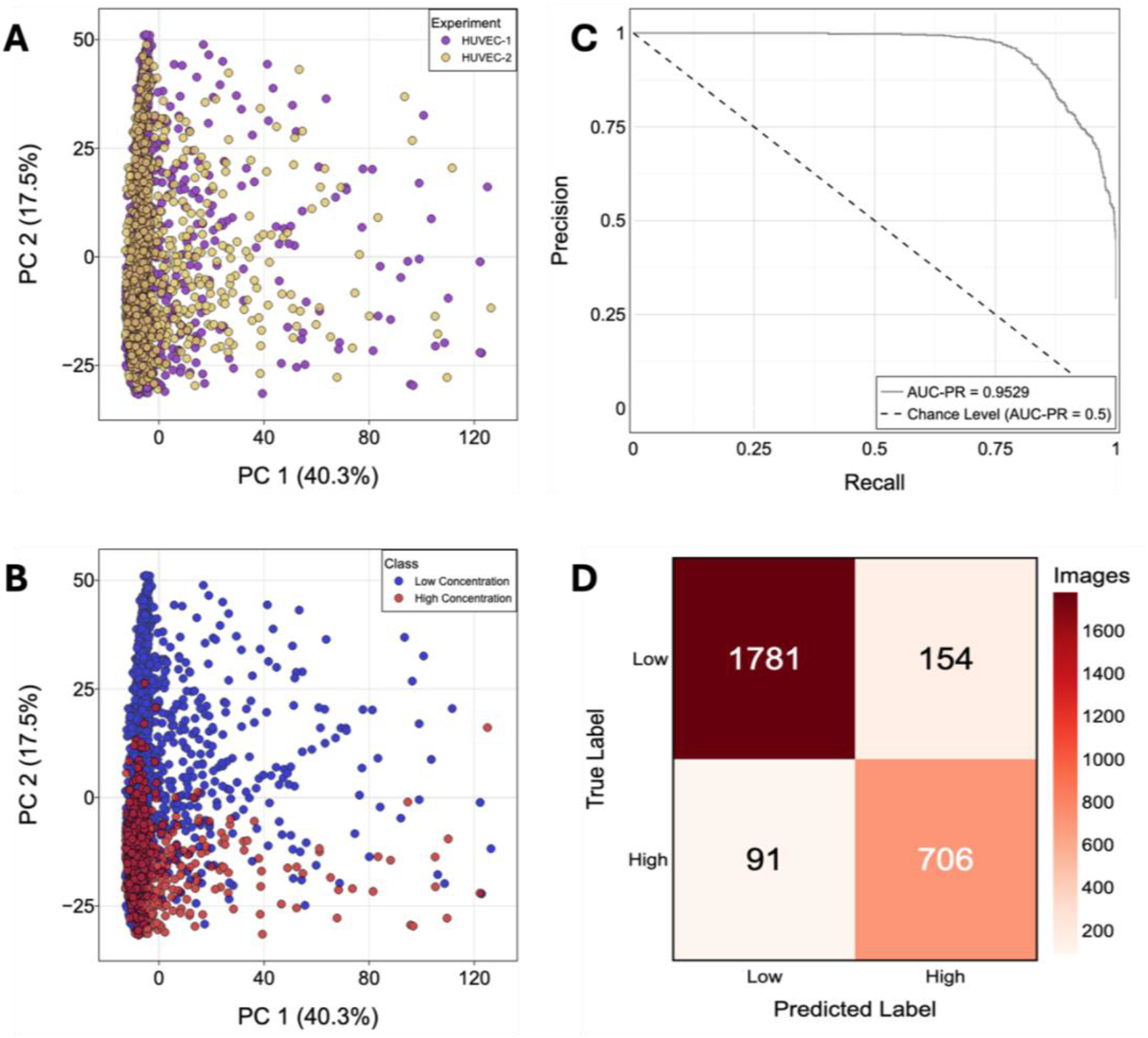
Performance of a binary classifier trained with DeepPaint. **A-B)** Principal component analysis (PCA) clustering of mean-centered and variance-scaled image embeddings. Each point represents an image in the test set. Images are colored by experiment **(A)** or binary class **(B)**. Percent variance captured on the first two principal components are denoted. Images with PC1 or PC2 scores outside of the 1^st^-99^th^ percentiles were omitted for visualization purposes. **C)** ROC curve for the binary classification model evaluated on our test set. The AUC is reported. **D)** Confusion matrix on our test set summarizing the number correctly and incorrectly classified images. The “Low” class includes untreated cells and cells treated with the three lowest tested concentrations of compound.

### Assessment of the Number of Training Images Required for High Model Performance

An important consideration when implementing a machine-learning model is the amount of training data required to make meaningful and statistically significant classifications during inference. To assess the amount of training data required to support the creation of a robust model, we plotted the average F1 score of our DeepPaint model as a function of the percentage of the total data used to train the model (**Figure *3***). Interestingly, a model retrained on 10% (∼1200 images) of the original training dataset was able to achieve an F1 score of 0.73, indicating a dataset of about 1200 images was sufficient for DeepPaint to generate a robust binary classifier.

**Figure 3.**
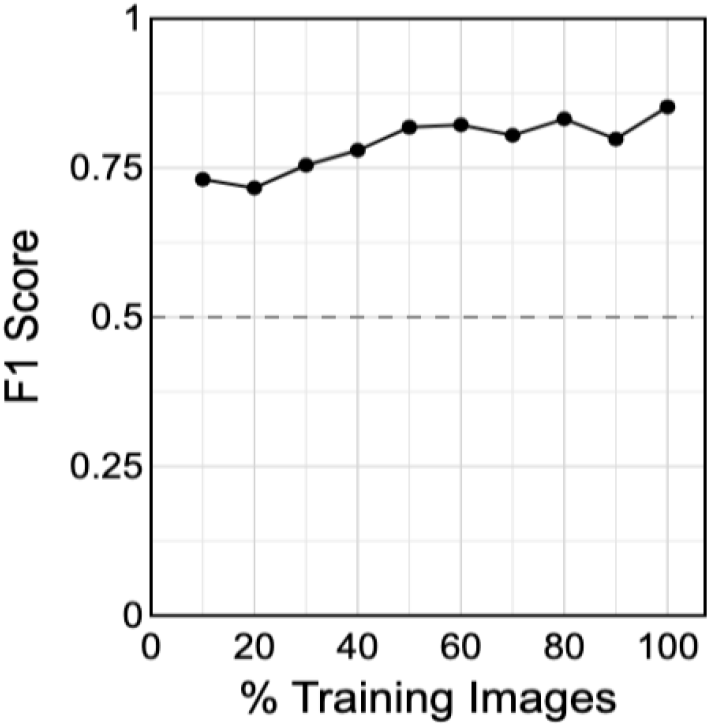
Test performance vs training dataset size. Images were randomly removed from the training dataset and the used to re-train a neural network to classify high and low dose images. The F1 score on the test dataset for each model is reported. Dashed line denotes expected performance of a random classifier.

### DeepPaint can be used to classify broad types of perturbations

Our next experiment aimed at applying our DeepPaint pipeline to a more realistic image classification problem. In this experiment we manually labeled our dataset into three classes: Cytokines/Growth Factors, Toxins, and Untreated (**Table *1***) to assess our model’s ability to cluster images by perturbant mechanism. We reasoned that because toxins function through very specific mechanisms, they should elicit a host cell response that is distinct from other soluble factors like cytokines and growth factors. When we retrained our model using this three-class problem we saw separation of our model output by class (**Figure *4*A, Error! Reference source not found**.) accompanied by high performance statistics (F1 score: 0.88, precision: 0.89, recall: 0.86) indicating that our model was able to differentiate between perturbant classes (**Figure 4D**). Close inspection of the confusion matrix (**Figure *4*E**) showed that many of the misclassifications by our model were of cytokines/growth factors as untreated cells. This was further highlighted by plots of the probability density distributions over each principal component (PC) for each class (**Figure *4*B-C**). We hypothesize that this is likely due to the modest phenotypic activities of various protein factors^25^ especially when delivered at low doses.

**Figure 4.**
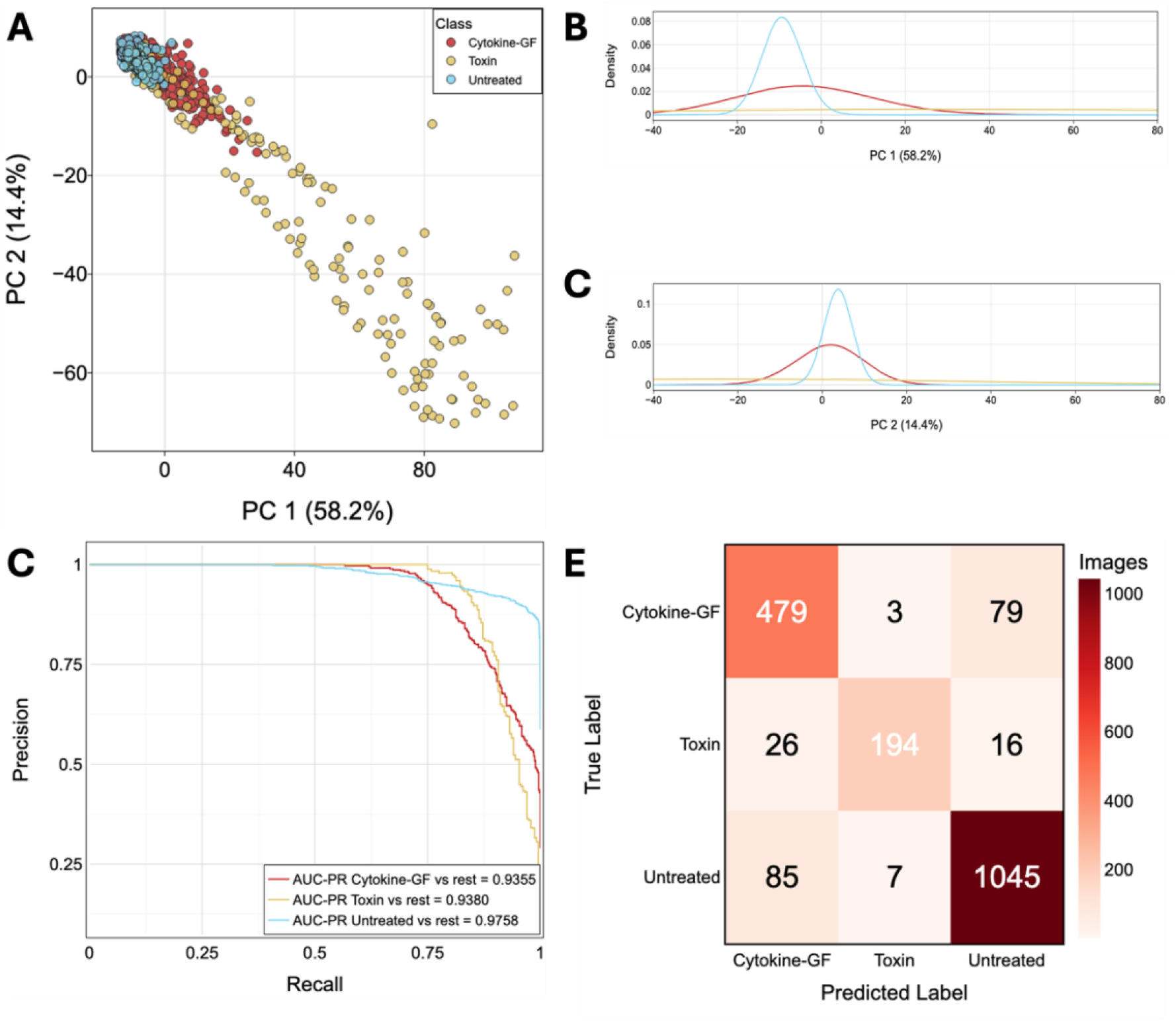
Performance of a multiclass classifier trained with DeepPaint. **A)** PCA clustering of mean-centered and variance-scaled image embeddings. Each point represents an image in the test set. Images are colored by broad treatment class (untreated = blue, cytokine-GF = pink, toxin = yellow). Percent variance on PC1 and PC2 are denoted. Density plots of **(B)** PC1 and **(C)** PC2 across each class are shown to help visualize the distribution of overlapping classes in the latent space. Images with PC1 or PC2 scores outside of the 1^st^-99^th^ percentiles were omitted from **(A-C)** for visualization purposes. **D)** ROC curves and AUC values for each class relative to all other classes. **E)** Confusion matrix on our test set summarizing the number of correctly and incorrectly classified images. Toxin and Cytokine-GF classes contain only high doses of each treatment.

## Conclusions

Here we presented DeepPaint, an open source, Python-based DenseNet-161 framework designed for Cell Painting image classification problems. DeepPaint integrates upstream image normalization and data augmentation pipelines to minimize data variability with downstream training and inference workflows for the classification of Cell Painting images without the need for image segmentation and *a priori* measurement designation. DeepPaint is a powerful framework for high-throughput screening of Cell Painting images (utilizing the network in an end-to-end fashion to classify multi-channel images) and discovery through analysis of embedded feature vectors. Given the ability to work directly on images, this approach greatly reduces the need for segmented cells and lowers the burden for human labeling. Our approach allows for the rapid retraining of the DenseNet for classification of new experimental data and rapid screening of new samples. This approach could also be combined with traditional analysis (*e*.*g*., CellProfiler) to explore the details of cell phenoprints in cases of special interest.

While DeepPaint is able to generate networks which can classify cell painting images with high-accuracy, models built on one dataset are not guaranteed to generalize well to new image sets. Alternative approaches which apply Weakly Supervised Learning frameworks to traditional cell-level and segmentation-based analyses (*e*.*g*., DeepProfiler^19^) are able to infer confounding factors and treatment outcomes from a CNN embedding in a potentially more generalizable way, but still require the need for single cell segmentation. Therefore, future efforts should aim to apply Weakly Supervised Learning frameworks with whole cell painting images to increase both generalizability and computational efficiency.

