## Supplementary figures and images for "DeepPaint: A deep-learning package for Cell Painting Image Classification"

### Supplementary Figure 1

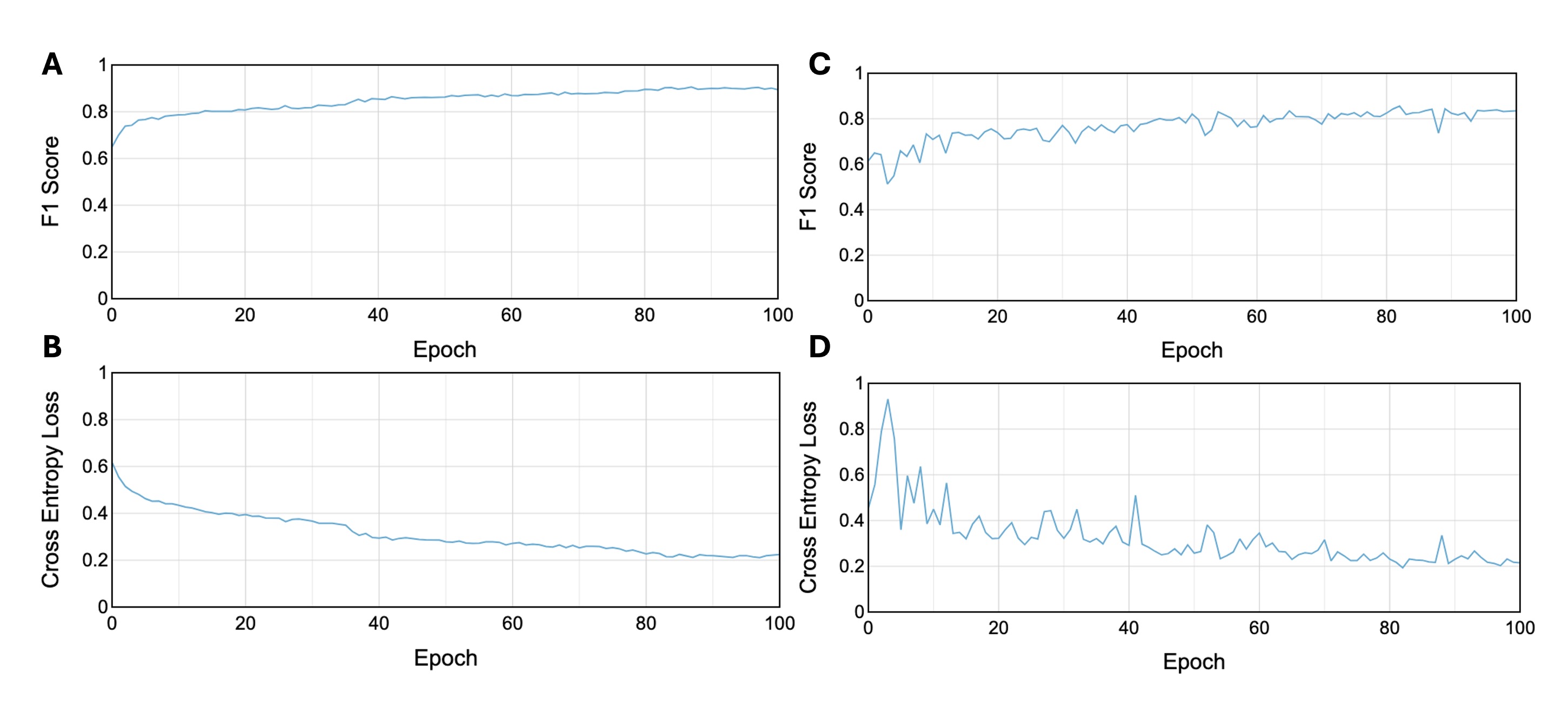

### Supplementary Figure 2

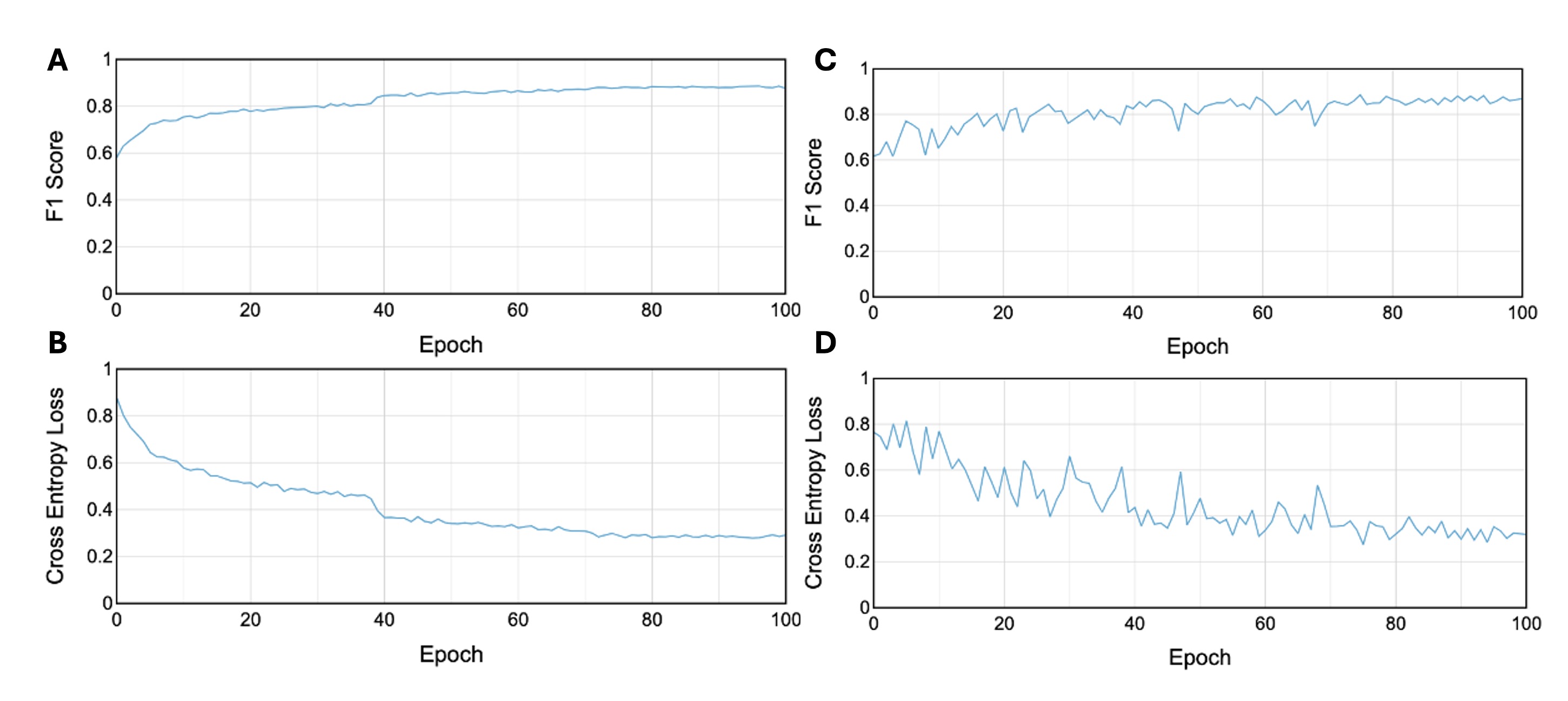
